# Expansion microscopy reveals unique ultrastructural features of pathogenic budding yeast species

**DOI:** 10.1101/2024.02.20.581313

**Authors:** Md. Hashim Reza, Srijana Dutta, Rohit Goyal, Hiral Shah, Gautam Dey, Kaustuv Sanyal

**Author notes:** These authors contributed equally: Rohit Goyal, Hiral Shah.

## Abstract

*Candida albicans* is the most prevalent fungal pathogen isolated from patients with candidemia. As is the case for many other fungi, the complex life cycle of *C. albicans* has been challenging to study with high-resolution microscopy techniques due to its small size. We employed ultrastructure expansion microscopy (U-ExM) to directly visualise sub-cellular structures at high resolution in the *C. albicans* yeast and during its transition to hyphal growth. NHS-ester pan-labelling in combination with immunofluorescence (IF) provided the first comprehensive map of nucleolar and mitochondrial dynamics through the *C. albicans* cell cycle. Analysis of microtubules (MTs) and spindle pole bodies (SPBs) stained with marker proteins suggests that contrary to the pole-to-pole arrangement observed in *Saccharomyces cerevisiae, C. albicans* yeast cells display a unique side-by-side arrangement of SPBs with a short mitotic spindle and longer astral MTs (aMTs) at the pre-anaphase stage. Modifications to the established U-ExM protocol enabled the expansion of several medically relevant human fungal pathogens, revealing that the side-by-side SPB configuration is a plausible conserved feature shared by many fungal species. We highlight the power of U-ExM to investigate sub-cellular organisation and organellar dynamics at high resolution and low cost in poorly studied, medically relevant microbial pathogens.

## Introduction

Since the first documented use of lenses to discover microbial life forms (Porter, 1976), various advancements have been brought about to improve the resolution of imaging. Conventional fluorescence microscopy is limited by low spatial resolution due to the diffraction limit of light that ranges from 200-300 nm laterally. Additionally, imaging cellular sub-compartments of fungi is limited due to the smaller-sized organelles, often beyond the diffraction limit of conventional fluorescence microscopes. The advent of super-resolution microscopy techniques, like structured illumination microscopy (SIM), photo-activated localisation microscopy (PALM) and stochastic optical reconstruction microscopy (STORM) have been able to achieve a resolution in the range of 50 - 120 nm (Betzig et al., 2006, Rust et al., 2006). The complexities associated with image acquisition and processing, coupled with the high cost of the microscopes, limit the throughput and benefits of super-resolution microscopy. The discovery of expansion microscopy (ExM), which relies on the isotropic physical expansion of biological samples rather than altered optics, enables super-resolution imaging using a diffraction-limited microscope (Chen et al., 2015). To date, the application of ExM to visualize ultrastructure in fungi is limited only to a few species including *Saccharomyces cerevisiae*, *Schizosaccharomyces pombe*, *Aspergillus fumigatus*, and *Ustilago maydis* (Chen et al., 2021, Götz et al., 2020, Hinterndorfer et al., 2022). This is largely due to a complex cell wall composition which prevents uniform expansion of the cell content in fungal species.

A common human microbiome resident, *Candida albicans* can transition from its otherwise commensal lifestyle to a pathogenic state (Mayer et al., 2013). *C. albicans* can switch between various morphotypes including yeast and hyphae. The yeast-hyphal transition is necessary for *C. albicans* pathogenicity, enabling tissue invasion and subsequent tissue damage during candidiasis (Sudbery et al., 2004, Lohse and Johnson, 2009). Additionally, plasticity with respect to ploidy, single nucleotide polymorphism (SNP), loss of heterozygosity (LOH), copy number variations (CNVs) and chromosomal instability (CIN) events, all make *C. albicans* a successful pathogen (Legrand et al., 2019, Selmecki et al., 2010). Of late, *C. albicans* has gained significant attention as a model organism for the study of nuclear division owing to attributes such as a dynamic genome, cryptic heterochromatin machinery (Sreekumar et al., 2019), and unique centromere properties (Legrand et al., 2019, Guin et al., 2020). While kinetochore proteins and their organisation have been studied in *C. albicans*, information regarding the spatial and molecular organisation of spindle pole bodies (SPBs) is largely lacking. SPBs, the functional equivalent of metazoan microtubule organising centres (MTOCs), nucleate nuclear and astral microtubules (aMTs) which segregate sister chromatids and position the spindle during the cell cycle, respectively (Markus et al., 2012, Palmer et al., 1992, Shaw et al., 1997, Sullivan and Huffaker, 1992). Positioning and alignment of the mitotic spindle along the polarity axis is vital for asymmetric cell division. The fungal kingdom displays remarkable diversity in the positioning of the mitotic spindle during the pre-anaphase stage of the cell cycle (Finley et al., 2008, Kopecka et al., 2001, Kozubowski et al., 2013, Maekawa et al., 2017, Markus et al., 2012, Martin et al., 2004, Mochizuki et al., 1987, Pereira et al., 2001, Winey and Bloom, 2012, Yamaguchi et al., 2009). In *S. cerevisiae*, the nucleus migrates to the bud neck with the mitotic spindle aligned to the bud axis at the pre-anaphase stage to achieve chromosomal division (Markus et al., 2012, Pereira and Yamashita, 2011, Winey and Bloom, 2012). However, both the nucleus and the mitotic spindle are positioned away from the bud neck in the pre-anaphase cells of C. *albicans* (Finley et al., 2008, Martin et al., 2004). This difference in SPB-dependent regulation of chromatid segregation further hints towards a distinctive feature of *C. albicans* cell biology that requires further exploration.

In this study, we established a working cell expansion protocol for *C. albicans* and succeeded in visualising sub-cellular structures in both planktonic yeast cells and hyphal germ tubes using NHS-ester pan-labelling (M’Saad and Bewersdorf, 2020) combined with immunofluorescence. We provide a characterisation of the mitotic cycle at ultrastructural resolution, revealing a unique configuration of SPBs in *C. albicans*. Finally, we demonstrate the applicability of U-ExM to six other important fungal pathogens.

## Results

### U-ExM reveals changes in the cellular ultrastructure during the yeast-hyphal transition in *C. albicans*

In most organisms with a cell wall, the nanoscale isotropic expansion that is critical to the U-ExM technique relies heavily on a post-fixation strategy to evenly digest the wall. Therefore, we optimised the digestion of the cell wall in the human fungal pathogen *C. albicans*. Log-phase chemically fixed cells were digested with Zymolyase 20T in a buffer containing 1.2 M sorbitol to prevent cell lysis. Post-digestion, cells were subjected to anchoring, followed by gelation, denaturation, and expansion (**Fig. 1A**).

**Figure 1.**
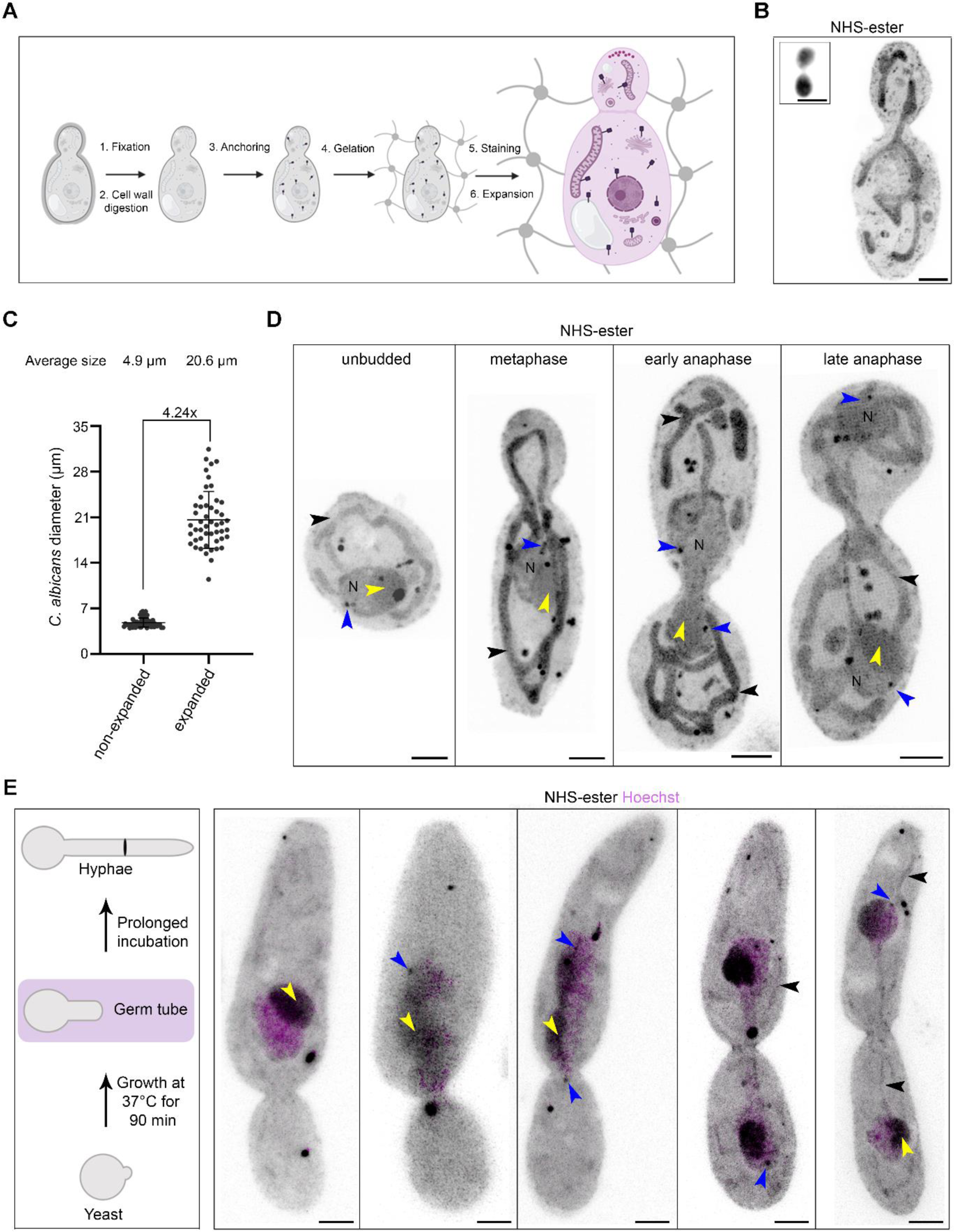
Pan-labelling of proteome displays sub-cellular organisation in expanded *Candida albicans* cells. (A) Schematic displaying different steps of ultrastructure expansion microscopy (U-ExM) protocol in *C. albicans*, which includes fixation (1), cell wall digestion (2), anchoring (3), gelation (4), staining (5), and expansion (6). (B) Representative confocal image of *C. albicans* cells post-expansion, pan-labelled with NHS-ester and displayed as maximum intensity projections. The inset shows a non-expanded cell. Scale bar 5 µm. (C) Scatter plot displaying the 4.24x expansion factor based on measurement of the diameter (longer axis) of unbudded cells of *C. albicans* before (*n*=54) and after (*n*=49) expansion. Error bars show mean±SD. (D) Representative images of *C. albicans* pan-labelled with NHS-ester showing sub-cellular organisation at various stages of the cell cycle. The black and blue arrowheads represent mitochondria and SPBs, respectively. The nucleus is marked with N and the nucleolus is marked with a yellow arrowhead. Scale bar 5 µm. (E) Schematic showing morphogenetic changes during the yeast-to-hyphal transition. Cells at the germ tube stage (magenta) were taken forward for U-ExM. Representative images of *C. albicans* pan-labelled with NHS-ester (grey) and co-stained with Hoechst (magenta) showing sub-cellular organisation at various stages of the cell cycle after hyphal induction. The black, blue, and yellow arrowheads represent mitochondria, SPBs, and nucleolus respectively. Scale bar 5 µm.

The gels were stained with an NHS-ester compound which non-specifically labels the proteome and enables visualisation of the protein density map of a cell (Kozubowski et al., 2013) (**Fig. 1B**). Next, we examined the degree of isotropic expansion and calculated the expansion factor. We measured the diameter of the *C. albicans* cell before (4.85 µm) and after expansion (20.6 µm) revealing that *C. albicans* could be expanded ∼4.24-fold (**Fig. 1C**). This is in line with the reported fold expansion for *S. cerevisiae* and *S. pombe* (Hinterndorfer et al., 2022). NHS-ester labelling highlighted specific organelles like mitochondria and nuclei. This experiment also enabled us to visualise the nucleolus as a region of higher protein density within the nucleus, and the SPBs as dark-stained punctate signals at the nuclear periphery and positioned away from the nucleolus (**Fig. 1D**).

*C. albicans* possesses a remarkable ability to switch between various morphological states, such as from unicellular yeast to hyphae, which are critical for virulence (Sudbery et al., 2004). We, therefore, sought to find if *C. albicans* hyphal cells can also be expanded and whether these cells display any structural variations from the yeast form. After hyphal induction by the addition of fetal bovine serum, the cells were fixed, digested, and expanded, as explained above. Pan-labelling revealed similar sub-cellular structures (nucleus, nucleolus, mitochondria, and SPBs) in hyphae as seen in the budding yeast (**Fig. 1E**). We observed a striking difference in both the number and shape of mitochondria in hyphae compared to yeast cells (**Fig. 1E**). By combining NHS-ester with Hoechst staining, we could capture the process of nuclear migration to the germ tube, elongated nuclei, nuclei connected by a mitotic bridge and finally segregated into two cells (**Fig. 1E**). We conclude that *C. albicans* can be fully expanded using U-ExM and that pan-labelling enables the identification of various sub-cellular structures and stages of cell division both in yeast and hyphal cells.

### Analysis of U-ExM images suggests organellar segregation patterns during the cell cycle are evolutionarily conserved

The cell cycle-dependent morphology of the mitochondrial network plays a central role in the growth and fitness of organisms by influencing metabolism and regulating various signalling cascades (Giacomello et al., 2020). Having seen a mitochondrial-like tubular network upon pan-labelling, we confirmed if these organelles were indeed mitochondria. For this, we co-stained *C. albicans* cells with the NHS-ester 405 and Bodipy^TR^ Ceramide. Bodipy^TR^ selectively stains lipid-rich organelles like the Golgi complex and mitochondria (Adisa et al., 2003). Indeed, co-staining confirmed the dense tubular network as mitochondria (**Fig. 2A**).

**Figure 2.**
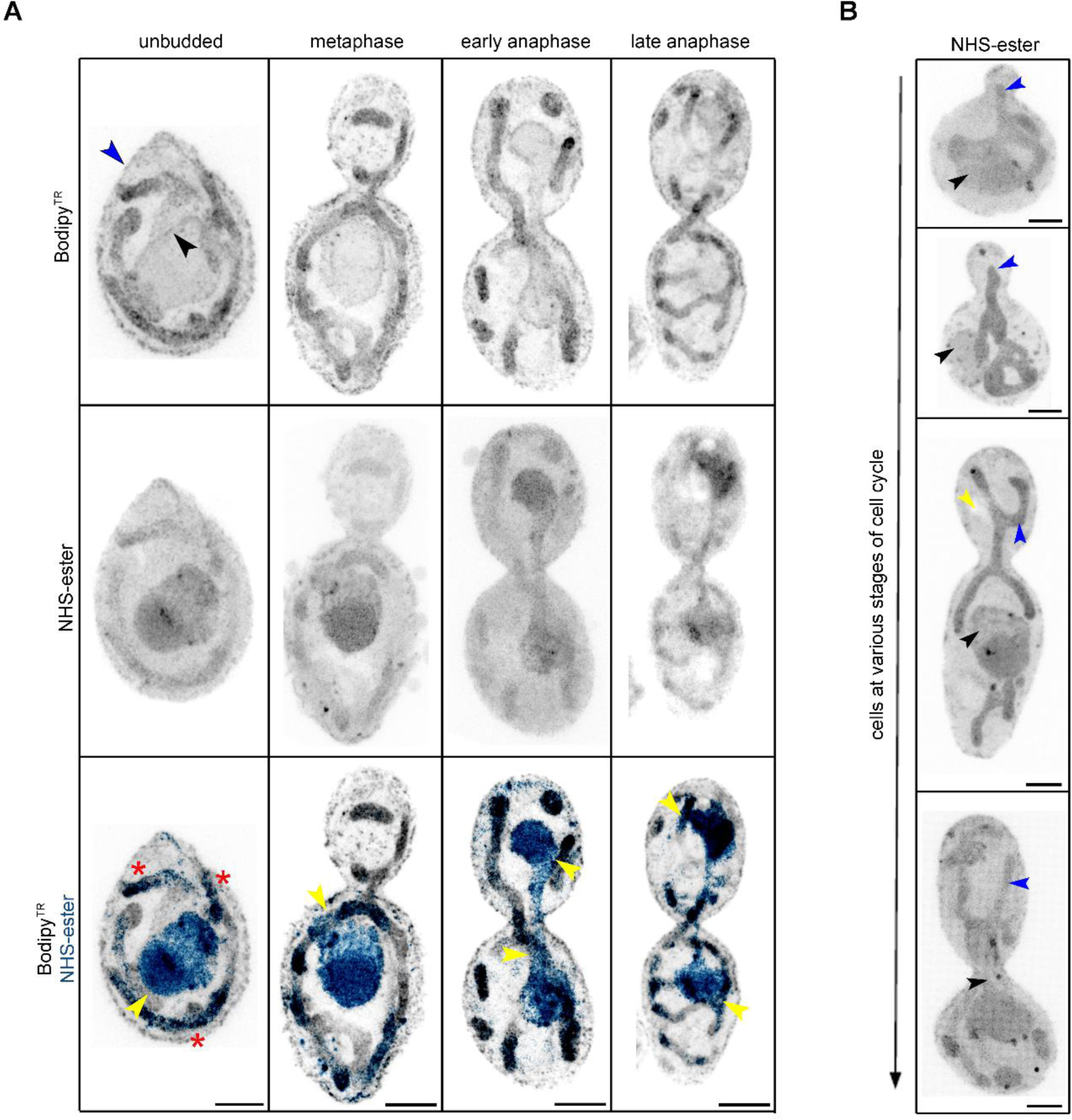
The tubular mitochondrial network segregates into daughter cells before nuclear segregation in *C. albicans*. (A) Maximum intensity projection of *C. albicans* cells co-stained with Bodipy^TR^ (grey) and NHS-ester (blue), at various stages of the cell cycle. The blue and black arrowheads mark the cell and nuclear membrane, respectively. The yellow arrowheads mark regions of close association of mitochondria with the nucleus. Red asterisks mark the mitochondria labelled with both Bodipy^TR^ and NHS-ester. Scale bar 5 µm. (B) Maximum intensity projection of *C. albicans* cells stained with NHS-ester (grey), during the cell cycle displays segregation of mitochondria (blue arrowheads) before nuclear (black arrowheads) segregation between the two cells. The yellow arrowhead marks the unlabelled vacuole in the cell. Scale bar 5 µm.

*C. albicans* displayed a tubular mitochondrial network at all stages of the cell cycle (**Fig. 2A**). A closer examination of U-ExM images also suggested a close association of mitochondria with the nucleus during the cell cycle (**Figs 1D, 2A**), hinting towards a likely crosstalk between these two organelles. Recently, a preferred order of organelle inheritance was shown in *S. cerevisiae*, wherein mitochondria are inherited before the migration of the nucleus into the daughter bud (Li et al., 2021). The analysis of the mitochondrial network and unstained vacuole during the cell cycle suggested that like *S. cerevisiae*, *C. albicans* cells consistently inherit both mitochondria and vacuoles before the migration of the nucleus during the cell cycle (**Fig. 2B**).

The NHS-ester pan-labelling also enabled us to focus closely on nuclear structures. We found a strong NHS-ester staining within the nucleus, which corresponded to the nucleolus (**Fig. 3A**). This was evidenced by Hoechst staining, which is mostly excluded from the nucleolus and stains chromatin (**Fig. 3A**). We also noticed a darkly stained region within the nucleolus of unknown identity (**Fig. 3B**). *C. albicans* did not exhibit a typical crescent-shaped nucleolus during interphase (**Fig. 3A**) as reported for *S. cerevisiae* (Girke and Seufert, 2019).

**Figure 3.**
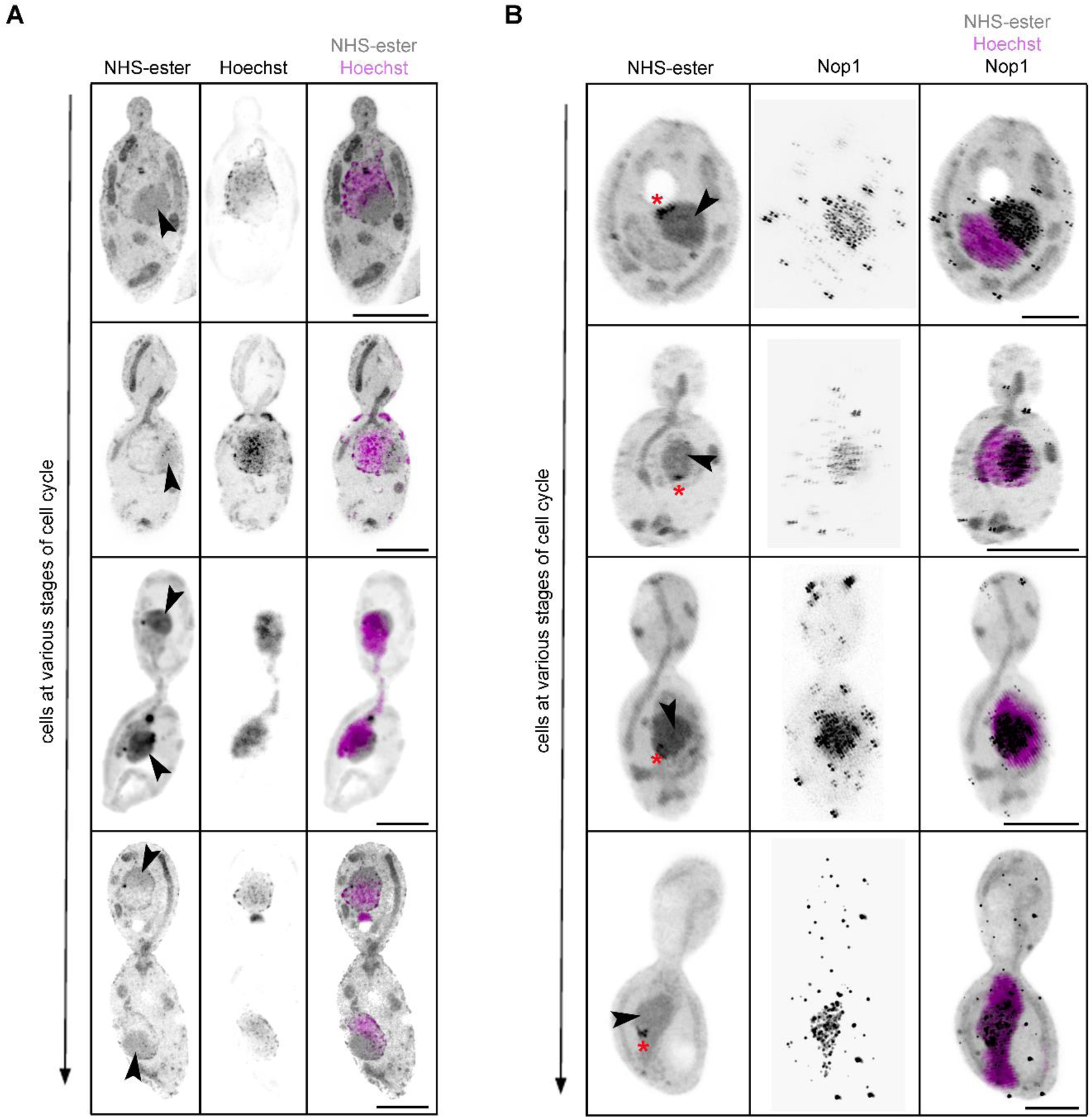
Nucleolar segregation during the cell cycle in *C. albicans*. (A) Maximum intensity projection of *C. albicans* cells co-stained with Hoechst (magenta) and NHS-ester (grey) during cell division. The black arrowheads mark the nucleolus. Scale bar 10 µm. (B) Maximum intensity projection of *C. albicans* cells co-stained with NHS-ester (grey), Hoechst (magenta) and Nop1 Abs (black) at various stages of the cell cycle. Scale bar 10 µm. The black arrowheads mark the nucleolus. The red asterisks represent a darker stained region within the nucleolus. Scale bar 10 µm.

Co-staining with NHS-ester and Hoechst helped us observe nucleolar segregation during the cell cycle in *C. albicans*. The nucleolus remained closely associated with the Hoechst-stained chromatin mass and segregated alongside bulk chromatin (**Fig. 3A**). This resembles nucleolar segregation dynamics seen in *S. cerevisiae* and in hyphal-induced *C. albicans* cells (Finley and Berman, 2005, Girke and Seufert, 2019, Granot and Snyder, 1991). We also validated the dark-stained region to be nucleolus by staining Nop1. Nop1 is a component of the small subunit processome complex and is required for the processing of pre-18s rRNA and localises to the nucleolus. Indeed, anti-Nop1 antibodies co-localised with the higher NHS ester-stained region within the nucleus during the cell cycle (**Fig. 3B**). Importantly, within the nucleolus, anti-Nop1 immunostaining revealed regions of higher and lower fluorescent intensities (**Fig. 3B**), reflecting differential intensities of Nop1. Taken together, NHS-ester pan-labelling provides an expansive view of the cellular landscape at various stages of the cell cycle in *C. albicans*.

### U-ExM provides insight into the organisation, assembly and inheritance of SPBs in *C. albicans*

SPBs nucleate microtubules, regulating nuclear positioning and spindle alignment during cell division. While the role of SPBs during the cell cycle is well known for the model yeasts *S. cerevisiae* and *S. pombe*, the organisation, assembly, and inheritance of SPBs are poorly understood in pathogenic fungi like *C. albicans*. Densely packed with proteins, SPBs in most species tend to be visible as a bright punctate structure in NHS-ester labelling (Shah et al., 2023, M’Saad and Bewersdorf, 2020) and also tend to be positioned away from the nucleolus in *Cryptococcus neoformans*, *Exophiala dermatitidis* and *S. cerevisiae* (Yamaguchi et al., 2010, Yamaguchi et al., 2003, Jin et al., 2000, Yang et al., 1989). In line with this, we also observed a bright punctate signal positioned away from the nucleolus and co-localising with chromatin (Hoechst staining) in *C. albicans* (**Figs 1D, S1A**).

Scale bar 2.5 µm. (D) Cartoon showing the mitotic spindle (green) between the two SPBs (red) dividing the Hoechst-stained chromatin (blue) area into two unequal segments (minor and major) as visually observed in a 2D-projected image. Violin plot showing percent area covered by the minor segment of Hoechst-stained chromatin in *C. albicans* and *S. cerevisiae*. *n*> 70 cells. Statistical analysis was done by Unpaired *t*-test with Welch’s correction (****p<0.0001).

To validate the identity of bright punctate structures as SPBs, we tagged Spc110, an inner plaque component of the SPB, with GFP and carried out IF using anti-GFP antibodies after the expansion of cells. Co-staining of Spc110 with NHS-ester confirmed these structures as SPB throughout the cell cycle (**Fig. 4A**).

**Figure 4.**
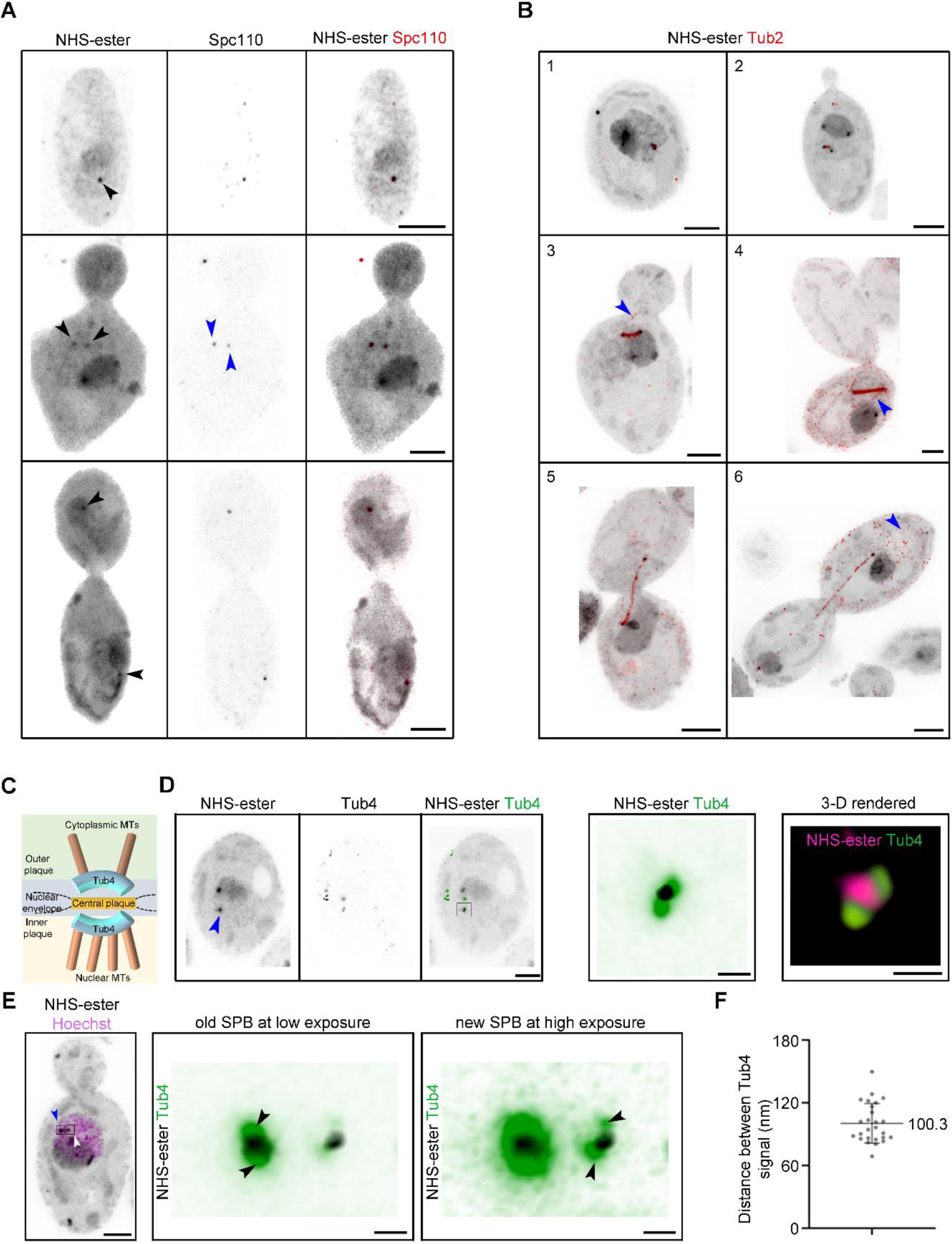
SPB organisation in *C. albicans*. (A) Maximum intensity projection of *C. albicans* cells co-stained with NHS-ester (grey) and anti-GFP (Spc110-GFP, red) through the cell cycle. The black and blue arrowheads mark the position of the SPBs stained with NHS and the position of the SPBs stained with anti-GFP, respectively. Scale bar 5 µm. (B) Maximum intensity projection of *C. albicans* cells co-stained with NHS-ester (grey) and anti-GFP (Tub2-GFP, red), at various stages of the cell cycle (1, 2, 3, 4, 5, and 6). The blue arrowheads mark the astral microtubules (aMTs). Scale bar 5 µm. (C) Schematic showing the spatial position of Tub4 at the SPB as reported from *S. cerevisiae*. (D) A representative maximum-intensity projection of *C. albicans* cells co-stained with NHS-ester (grey) and anti-GFP (Tub4-GFP, red) in unbudded cell. Scale bar 5 µm. Inset, a magnified region depicted by a black-bordered square and a 3-D rendered image of the magnified region is also shown. Scale bar 125 nm. (E) Maximum-intensity projection of *C. albicans* cells co-stained with NHS-ester (grey) and Hoechst (magenta) at the pre-anaphase stage of the cell cycle. Scale bar 5 µm. The blue and white arrowhead represents the old and new SPBs, based on signal intensity. Inset, a magnified region is shown for the two SPBs. The first inset highlights the Tub4 arrangement (black arrowheads) at the old SPB, visible at low-intensity exposure. The second inset highlights the Tub4 arrangement (black arrowheads) at the new SPB visible only at high-intensity exposure. Scale bar 125 nm. (F) Scatter plot displaying the distance between two Tub4 fluorescent signals rescaled after expansion (N=2, *n*=26). The error bar shows mean ± SD.

Importantly, immunostaining revealed an asymmetry between both SPBs, which was evident till the cells progressed into anaphase (**Fig. 4A**). This differential staining was also observed upon NHS-ester labelling, going on to show that similar to *S. cerevisiae* (Liakopoulos et al., 2003), the old SPB in *C. albicans* has a higher protein density. We also validated these structures to be the MT nucleation centres by immunostaining tubulin in a strain carrying Tub2-GFP (**Fig. 4B**). U-ExM revealed the structural changes in the mitotic spindle during the cell cycle, with the spindle being compact at early cell cycle stages, evident from intense staining (stages 1-4, **Fig. 4B**). As the cells enter into anaphase, the mitotic spindle shows low staining, due to the presence of fewer kinetochore MTs (stages 5-6, **Fig. 4B**), as reported for *S. cerevisiae* (Winey and O’Toole, 2001).

The SPB structure and its duplication during the cell cycle are well-studied in two model yeasts, *S. cerevisiae* and *S. pombe* (Cavanaugh and Jaspersen, 2017). The SPB is divided into inner, central, and outer plaque in *S. cerevisiae*. The central plaque anchors the outer and inner plaques which nucleate astral/cytoplasmic and nuclear microtubules, respectively. One of the applications of isotropic expansion is the decrowding of the intracellular space which provides the tool to study the effective spatial resolution of proteins. To investigate the effective resolution of two plaques of SPB in *C. albicans*, we resorted to γ-tubulin homolog, Tub4, positioned on both inner and outer plaques (**Fig. 4C**). GFP-tagged Tub4-expressing *C. albicans* cells were expanded and probed with anti-GFP antibodies. Using Airyscan imaging, we could obtain two Tub4 fluorescence signals, representing the inner and outer plaques in unbudded cells (**Fig. 4D**). We were also able to detect four Tub4 dots, two from each SPB, post-SPB duplication (**Fig. 4E**). The more intense old SPB, evident from NHS-ester labelling, showed symmetric Tub4 signals between the inner and outer plaque. However, we observed a difference in the signal intensity between the two Tub4 signals for the less intense new SPB (**Fig. 4E**). We estimated the distance between the two Tub4 signals after 2D-projection and found them to be separated by 100.3±5.3 nm, after rescaling with the expansion factor. Altogether, using U-ExM, we could study SPB asymmetry and estimate the distance between the outer and inner plaque by resolving Tub4 fluorescent signals in *C. albicans*.

Separation of the duplicated SPBs, followed by their movement to the diametrically opposite sides of the nuclear envelope, is a prerequisite for the formation of a bipolar mitotic spindle in *S. cerevisiae* (Cavanaugh and Jaspersen, 2017). To study the SPB separation dynamics in *C. albicans*, we tagged the spindle with Tub2-GFP and SPBs with Tub4-mCherry. For comparison, spindle and SPBs were tagged with GFP-Tub1 and Spc42-mCherry, respectively in *S. cerevisiae*. We looked at the distribution of the pre-anaphase spindle length with respect to the budding index. We found that the spindle in *C. albicans* was restricted to a length of <1 µm when compared to the 1-1.5 µm spindle in *S. cerevisiae* (**Fig. S1B**), resulting in unequal partitioning of the 2D-projected chromatin-covered nuclear area (major and minor segments) (**Fig. S1C**). To validate this, we measured the proportion of chromatin covered by the minor segment (see materials and methods) in pre-anaphase cells. In *C. albicans*, the minor segment constitutes 23% of the chromatin area, whereas in *S. cerevisiae* we observed a 32% coverage (**Fig. S1D**), suggesting a side-by-side arrangement of SPBs in *C. albicans* as opposed to a pole-to-pole arrangement in *S. cerevisiae*. The U-ExM images with tubulin staining also suggested very long aMTs positioned all along the cell cortex (**Supplementary Movie 1**). *C. albicans* exhibits free cytoplasmic MTs (cMTs) and their presence is cell cycle-dependent (Finley and Berman, 2005, Lin et al., 2016). The appearance of free cMTs mirrors the depolymerization of MTs during telophase (**Fig. S2A**). These free cMTs persist till SPBs are duplicated and decline as cells enter metaphase (**Fig. S2A**). In summary, *C. albicans* shows an atypical arrangement of duplicated SPBs with shorter pre-anaphase spindles along with the presence of long aMTs and free cMTs, suggesting different MT dynamics and regulation compared to *S. cerevisiae*.

### U-ExM as a broadly applicable tool to investigate ultrastructure in human fungal pathogens

We extended the optimised U-ExM protocol of *C. albicans* to several human fungal pathogens belonging to diverged fungal lineages: the CUG-Ser1 clade and the WGD lineage of Ascomycota, and a species that belongs to Basidiomyota. Modifications in the *C. albicans* ExM protocol for each of these species are included in materials and methods. For most of these species, increasing the incubation time from 45 min to 60 min for cell wall digestion facilitated near-complete expansion, except for *Candida auris*, *Candida tropicalis*, and *C. neoformans*. *C. neoformans* required an additional Triton X-100 treatment before the cell wall digestion for increased efficiency.

We demonstrate that *Candida dubliniensis* and *Candida parapsilosis,* two CUG clade species, related to *C. albicans*, could be expanded 4.21 and 3.58-fold, respectively (**Figs 5A, B**). *C. tropicalis* and *C, auris*, also belonging to the CUG-Ser1 clade showed an expansion factor of 3.03 and 2.92, respectively (**Figs 5C, D**). While, the ascomycete, *Nakaseomyces glabratus* which belongs to the WGD clade could be expanded by 3.96-fold, the basidiomycete *C. neoformans* showed an expansion factor of 2.48-fold only (**Figs 5E, F**). We could observe various sub-cellular structures like nuclei, nucleolus, SPBs, and mitochondria in the expanded human fungal pathogens. Together, we demonstrate a successful optimisation of the expansion of *C. dubliniensis*, *C. parapsilosis*, and *N. glabratus* (**Fig. 5G**). The differences in several sub-cellular structures were evident across the species. Mitochondria in *C. dubliniensis* and *N. glabratus* were tubular, while *C. tropicalis* displayed both tubular and fragmented mitochondrial networks (**Figs 5A-F**). Having seen a side-by-side arrangement of SPBs in *C. albicans*, we were curious to know the SPB arrangements in *C. dubliniensis* and *N. glabratus.* Remarkably, we observed a side-by-side arrangement of SPBs in both these species (**Figs 5H, I**). Thus, U-ExM followed by pan-labelling identified the conservation in side-by-side SPB arrangements in these two species.

**Figure 5.**
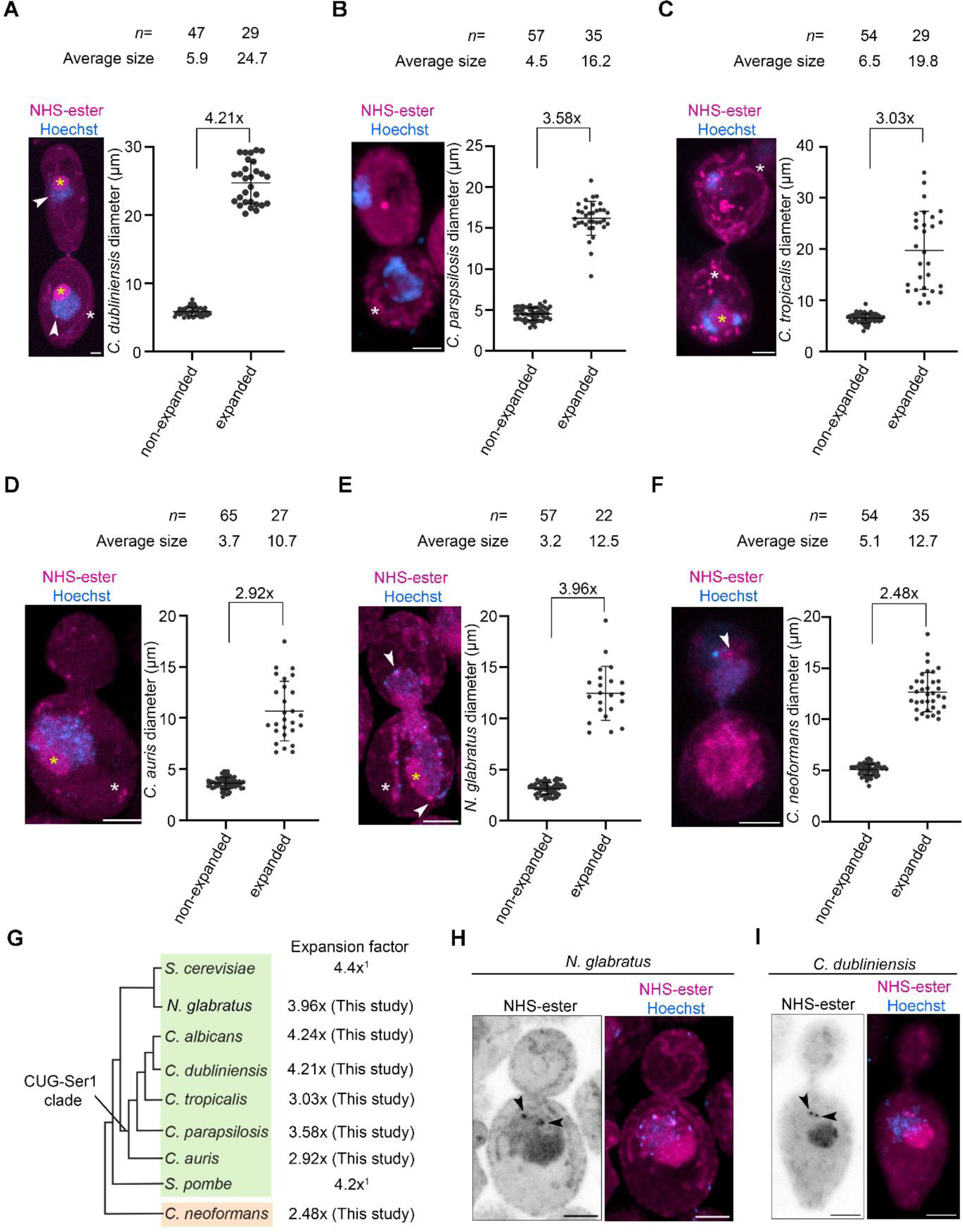
Expansion followed by pan-labelling reveals sub-cellular organisation in human fungal pathogens. (A-F) Confocal images of human fungal pathogens post-expansion, co-stained with NHS-ester (magenta) and Hoechst (blue) and displayed as maximum intensity projection (*left*). Corresponding scatter plot (*right*) displaying the expansion factor based on measurement of the diameter of unbudded cells for these species before and after expansion. Error bars show mean±SD. The white arrowheads represent SPBs. The nucleolus and mitochondria are labelled with yellow and white asterisks, respectively. Scale bar 5 µm. (G) Cladogram showing the human fungal pathogens used in this study for U-ExM, Acomycota (green) and Basidiomycota (orange). The expansion factor for these fungal pathogens is mentioned along with the well-known budding and fission yeast, *S. cerevisiae* and *S. pombe* (1: (Hinterndorfer et al., 2022)), respectively. (H) Representative confocal image of *N. glabratus* post-expansion, pan-labelled with NHS-ester (magenta) and Hoechst (blue) and displayed as maximum intensity projection. The black arrowheads show sideward SPB arrangements (*n=*19, budding index= 0.33-0.69). (I) Representative confocal image of *C. dubliniensis* post-expansion, pan-labelled with NHS-ester (magenta) and Hoechst (blue) and displayed as maximum intensity projection. The black arrowheads show sideward SPB arrangements (*n=*12, budding index= 0.47-0.75). Scale bar 5 µm.

## Discussion

In this work, we optimised the U-ExM protocol in the human fungal pathogen *Candida albicans*, a model system to study cell division and host-pathogen interactions. Expansion together with pan-labelling of the proteome facilitated the monitoring of organellar segregation dynamics during cell division of *C. albicans* in both yeast and hyphal forms. With a 4-fold expansion, we could successfully resolve the inner and outer plaques of the SPBs. We extended this protocol to expand its application to some of the non-model medically relevant human fungal pathogens. We establish U-ExM as a powerful tool to study the cell biology of model and non-model fungal systems.

Dual dye staining revealed the tubular mitochondria migrating before the nucleus into the daughter cell in *C. albicans*, while the nucleolus moves in conjunction with chromatin during the cell cycle, as evident from immunostaining of Nop1. This highlights the compatibility of U-ExM with dual dye and immunostaining. Unlike the well-studied model organism *S. cerevisiae*, a non-crescent-shaped nucleolus was observed with seemingly large occupancy in the *C. albicans* nucleus. Important for ribosome biogenesis and regulation, a change in size, number and structure of nucleolus is often associated with various cellular metabolic states (McCann and Baserga, 2014). While nutrient restriction leads to a reduction in size (Matos-Perdomo and Machín, 2019), metabolically active cells have enlarged nucleolus (Weeks et al., 2019). Whether a difference in the lifestyle between pathogenic (*C. albicans*) and non-pathogenic (*S. cerevisiae*) organisms relates to a differential nucleolar shape and occupancy, is an area for further research.

U-ExM is an excellent tool for studying differential protein occupancy which becomes pronounced due to a dilution of fluorescence intensity upon expansion (Götz et al., 2020). This was evident for SPB proteins, Spc110 and Tub4, which revealed a difference in protein density between the old and new SPBs. The asymmetric Tub4 distribution was also apparent between the inner and outer plaque at the newly formed SPBs. Unlike *S. cerevisiae*, *C. albicans* does not show any noticeable asymmetric distribution of Tub4 between the two plaques during interphase and in the old SPBs, post-duplication by U-ExM. A shorter Tub4-to-Tub4 distance in *C. albicans* further hints towards an SPB organisation distinct from *S. cerevisiae* (Burns et al., 2015, Byers and Goetsch, 1974, Hinterndorfer et al., 2022). *C. albicans* displaying a side-by-side SPB arrangement in pre-anaphase unlike the pole-to-pole arrangement seen in *S. cerevisiae*, further hints towards a difference in SPB separation events. Thus, we could obtain a pronounced view of SPB dynamics and critically analyse the differences in SPB features in *C. albicans* using U-ExM, overcoming the limitations of conventional microscopy.

In this study, we show that the U-ExM protocol can be applied to six other human fungal pathogens, belonging to Ascomycota and Basidiomycota fungal phyla. The cell wall of *C. albicans* is known to have more β-1,6-glucans compared to *S. cerevisiae* (Brown and Gordon, 2005), which is reflected in the timing of cell wall digestion, with *C. albicans* requiring a longer time for complete cell wall digestion than *S. cerevisiae* (Hinterndorfer et al., 2022). In our study, a 4-fold expansion could not be achieved for *C. auris*, *C. tropicalis*, and *C. neoformans*. Composed of α-1,3-glucan, β-1,3 and β-1,6-glucan, chitin, chitosan, mannoproteins and GPI-anchored proteins (Garcia-Rubio et al., 2020), the two-layered cell wall of *C. neoformans* is further surrounded by an exopolysaccharide capsule (Garcia-Rubio et al., 2020). This vastly differs from the cell wall properties of *C. albicans* and related *Candida* species, explaining the reduction in the expansion factor observed. On the other hand, the *C. auris* and *C. tropicalis* isolates used in this study are resistant and tolerant to fluconazole (National Culture Collection of Pathogenic Fungi (nccpf.in)), respectively, which is implicated with increased levels of cell wall chitin (Shahi et al., 2022). This could be the likely reason for the inefficiency in achieving a 4-fold expansion despite *C. auris* and *C. tropicalis* being related species to *C. albicans*.

The unique side-by-side arrangement of SPBs observed in this study for *C. albicans* was also reflected by the NHS-labelled SPBs in *C. dubliniensis* and *N. glabratus*. This demonstrates the importance of U-ExM in the study of various cell biological processes in the absence or relative ease of techniques available for native tagging, live-cell microscopy, and standardised transformation protocols of various non-model organisms. Our analysis suggests that despite being a member of the WGD clade, *N. glabratus* SPBs do not follow segregation dynamics akin to *S. cerevisiae*. This divergence in SPB dynamics between WGD species calls for further studies as molecular details regarding the role of SPBs in asymmetric cell division, inheritance and evolution in fungal pathogens are still lacking.

## Materials and Methods

### Yeast strains and culture

All the strains used in this study are specified in Supplementary Table S1.

### Reagents used in the study

The following primary antibodies were used in this study: anti-Nop1 (Anti-Fibrillarin antibody [38F3] - Nucleolar Marker (ab4566)) used at 1:500, anti-GFP (mouse) (Roche, 11814460001) used at 1:500. The following secondary antibodies were used in this study: Alexa fluor 488 goat anti-mouse IgG (Invitrogen, A11001). The secondary antibodies were used at 1:500 dilution. The following NHS-ester dyes were used in the present study: Dylight^TM^ 405 NHS-ester (Thermo Fisher Scientific, 46400), Dylight^TM^ 594 NHS-ester (Thermo Fisher Scientific, 46412), and Alexa Fluor^TM^ 647 carboxylic acid, Succinimidyl ester (Thermo Fisher Scientific, A20006), all used at 1:500. Bodipy^TR^ Ceramide (Invitrogen D7540), Formaldehyde (Fischer Scientific, 24008), Acrylamide (Merck, A4058), N, N’-Methylenebisacrylamide (Merck, M1533), Sodium acrylate (Merck, 408220-256), Ammonium persulphate (APS) (HiMedia, MB003), TEMED (Merck, T7024), Hoechst 33342 (Sigma, B2261).

### Yeast culture, fixation, and cell wall digestion

Briefly, log-phase cells (approximate 1 optical density at 600 nm) were first fixed with 3.7% formaldehyde for 15 min at 30°C with intermittent shaking. 1 OD equivalent cells were taken forward for subsequent cell wall digestion. For *C. albicans*, C*. tropicalis*, *N. glabratus*, *C. dubliniensis* and *C. parapsilosis*, cells were washed twice with PEM buffer (100 mM PIPES, 1 mM EGTA, 1 mM MgSO_4_, pH to 9.0) at 5,000 rpm for 5 min and 15 min for *C. auris*, and once with PEM-S (1.2 M sorbitol in PEM) at 5,000 rpm for 5 min and 15 min for *C. auris*. The fixed cells resuspended in 100 µL of PEM-S buffer were enzymatically digested with a final concentration of 2.5 mg/mL Zymolyase 20T at 30°C for 45 min in the case of *C. albicans*, *C. dubliniensis*, and C*. tropicalis* and 1 h in case of *C. auris*, *C. parapsilosis* and *N. glabratus*. Cells were washed once with PEM-S buffer at 5,000 rpm for 5 min and 15 min for *C. auris* and the resuspended cells were proceeded for anchoring.

For *C. neoformans*, 1 OD equivalent cells were washed with PEM and PEM-S buffer as mentioned. Cells were then resuspended in 500 µL of PEM-S buffer with 0.2% Triton X-100 and incubated at 30°C for 30 min at 100 rpm. It was followed by digestion using 25 mg of Lysing enzyme (Sigma, L1412) dissolved in 500 µL of PEM-S buffer and incubated at 30°C for 6 h at 100 rpm. Digested cells were taken forward for anchoring.

### Yeast-to-hyphal induction

Briefly, log-phase *C. albicans* cells (approximate 1 optical density at 600 nm) were added to pre-warmed media (at 37°C) containing 9 mL of YPD+uridine (10 µg/mL) and 1 mL of Fetal bovine serum (Thermo Fisher Scientific, 10270106). The cells were grown at 37°C for 90 min at 180 rpm for the induction of germ tube formation. The cells were fixed and proceeded for U-ExM as described earlier. Post-germ tube formation, cells were pelleted down at 5,000 rpm for 15 min for every step involving centrifugation.

### Ultrastructure Expansion microscopy (U-ExM)

U-ExM was performed as previously described (Hinterndorfer et al., 2022), with a few modifications. The digested cells were kept for anchoring in acrylamide (AA)/formaldehyde (FA) (1% AA, 0.7% FA diluted in 1x PBS) overnight at 37°C, 12 rpm on a Rotaspin. The next morning, a 6 mm coverslip was coated with poly-L-lysine (Sigma, P8920) for 1 h at room temperature. The anchored cells were then allowed to attach to the Poly-L-lysine coated coverslip for 1 h. Gelation was performed on ice using a cocktail of monomer solution (19% (wt/v) sodium acrylate, 10% (v/v) acrylamide, 0.1% (v/v) N, N’-methylenebisacrylamide in PBS), TEMED (0.5% v/v) and APS (0.5% v/v). The cells were incubated for 10 min on ice. The gel was kept for polymerization for 1 h at 37°C in a moist chamber. Next, the gel was transferred to 1 mL denaturation buffer (50 mM Tris pH 9.0, 200 mM NaCl, 200 mM SDS, pH to 9.0) and incubated at 95°C for 1 h 30 min at 300 rpm. After denaturation, the gel was expanded with three subsequent washes with water, 15 min each. The gel diameter was measured after expansion to determine the expansion factor. The gels expanded in the range of 3.7-4.4 fold. The gel was shrunk with three washes of 1x PBS for 10 min each. Pan-labelling for U-ExM was done using Dylight^TM^ 405 NHS-ester/ DyLight^TM^ 594 NHS-ester/ Alexa Fluor^TM^ 647 carboxylic acid in 1x PBS overnight at 4°C.

### Immunofluorescence staining

For Nop1 and GFP immunostaining, the gel was stained using anti-Nop1 and anti-GFP as the primary antibody at 1:500 and incubated overnight at 4°C. The gel was washed thrice with PBS with 0.1% Tween 20 for 30 min at room temperature. Gel was then incubated with goat anti-mouse-IgG coupled to Alexa Fluor 488 secondary antibody at 1:500 and incubated at 37°C for 3 h in the dark. The antibody dilutions were prepared in 3% BSA in 1x PBS with 0.1% Tween 20. The gel was washed thrice with PBS with 0.1% Tween 20 for 30 min at room temperature. The gel was expanded with three subsequent washes with water before imaging.

### Sample mounting and imaging

For microscopy, Poly-L-lysine coated Ibidi chamber slides (2 well – Ibidi 80287) or MatTek glass bottom dishes (P35G-0-14-C) or Cellvis (2 Chambered Coverglass system-C2-1.5H-N) were used. Gels were cut to an appropriate size to fit the glass bottom chambers and were overlaid with water to prevent drying or any shrinkage during imaging. The gels were imaged using the Zeiss LSM980 Airyfast confocal microscope using a Plan-Apochromat 63x/1.4 Oil DIC M2pb7 objective or LSM880 Airyfast confocal microscope using a Plan-Apochromat 63x/1.4 Oil DIC M27 or 100x/1.4 Oil DIC M27 objectives.

For Figures 1E, 4 A-F, and 5 A-H, the gels were imaged with a Zeiss LSM880 AiryFast confocal microscope using a 63x oil-immersion objective (NA 1.42) at a step size of 0.3 µm.

For U-ExM images, scale bars have not been rescaled for the gel expansion factor.

### Quantification of Tub4 distance in expanded *C. albicans* cells

The Tub4 distance between the inner and outer plaque of the SPB was quantified as the distance between the maximum intensities of both the signals which corresponded to the two plaques. The Tub4-to-Tub4 distance was normalized with the expansion factor.

### Quantification of SPB-to-SPB distance and nuclear partitioning by the spindle

The budding index for non-expanded cells was calculated by measuring the ratio of the diameter of the daughter and mother bud. The SPB-to-SPB distance was measured as the distance between the centre of two Tub4-mCherry (*C. albicans*) and Spc42-mCherry (*S. cerevisiae*) signals. Both budding indices and SPB-to-SPB distances were calculated using a straight-line selection tool from Fiji. SPB-to-SPB distance and budding index were calculated for cells having two SPB signals thus excluding the unbudded cells and small-budded cells with single SPB puncta. To understand the proximal arrangement of SPBs on the nucleus, the Hoechst-stained nucleus was divided into two segments by a straight line covering the spindle and passing through the SPBs. The area of the minor segment of the Hoechst-stained region and the whole region was calculated using the Freehand selection tool of Fiji and the percentage of area covered by the minor segment was used as data points.

### Statistical analysis

Statistical analysis was done using GraphPad Prism 8.4.0 software. Student’s Unpaired *t*-test followed by Welch’s correction for comparing two groups.

## Acknowledgements

We thank the members of GD and KS lab for the valuable discussions and comments. We thank the European Molecular Biology Laboratory (EMBL) Advanced Light Microscopy Facility (ALMF) for the technical input. We thank B. Suma and S. Patil at the imaging facility at JNCASR.

## Author contributions

Conceptualization: MHR, GD, KS; Methodology: HS, MHR, SD, RG; Validation: MHR, SD, RG, HS, GD, KS; Formal analysis: MHR, SD, RG, HS; Writing-original draft: MHR, SD, RG, HS, GD, KS; Writing-review & editing: MHR, SD, RG, HS, GD, KS; Visualization: MHR, SD, RG, HS; Supervision: GD, KS; Funding acquisition: MHR, GD, KS.

## Data availability

All data on the results can be found within the article and supporting information.

---

**Figure S1.**
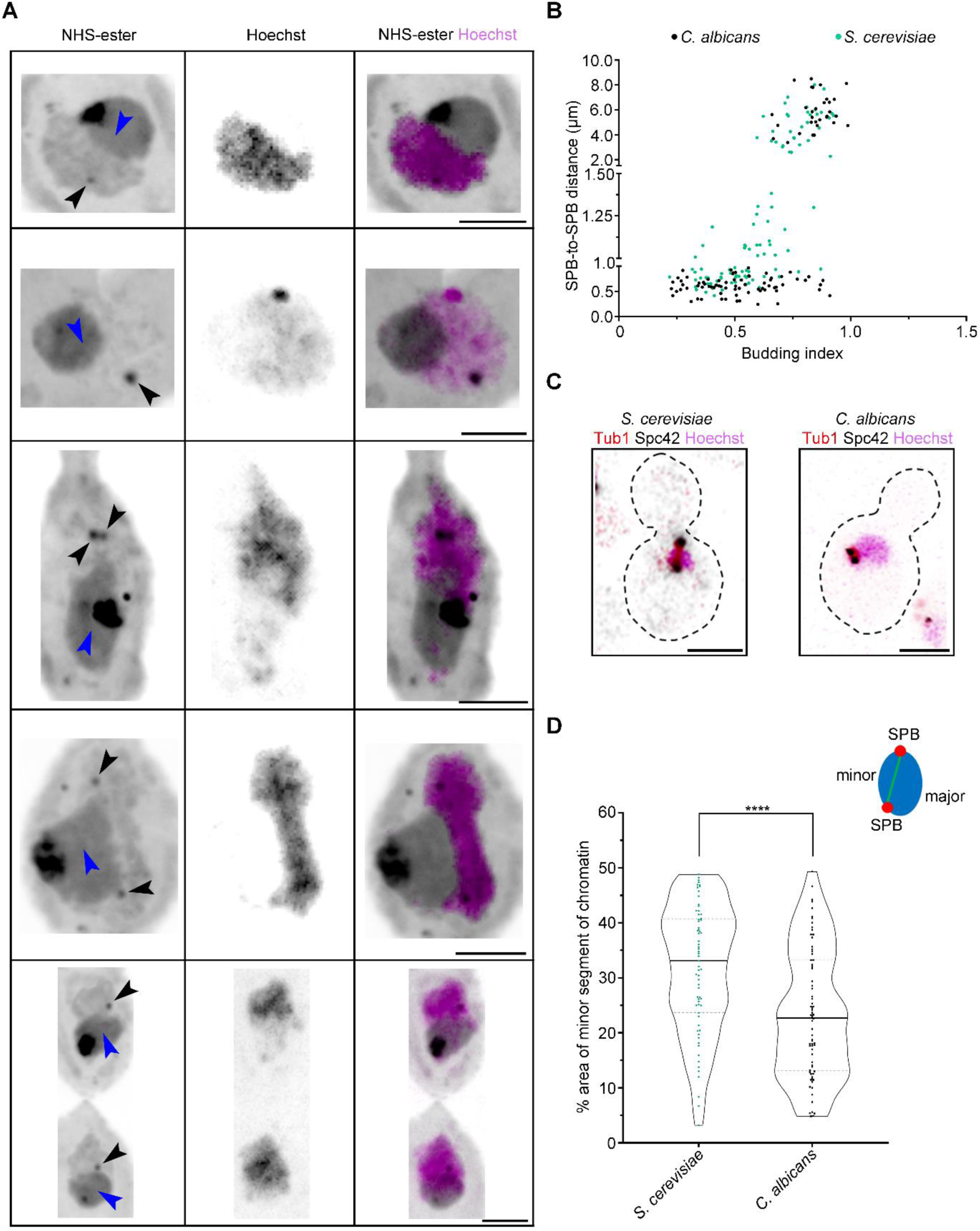
SPB positioning during cell division in *C. albicans*. (A) Zoomed image showing the nucleus, co-stained with NHS-ester (grey) and Hoechst (magenta) during the cell cycle. The black arrowheads mark the position of the spindle pole bodies (SPBs) away from the nucleolus (blue arrowheads). Scale bar 5 µm. (B) Scatter plot displaying SPB-to-SPB distance with respect to the budding index in *C. albicans* (black) and *S. cerevisiae* (green). SPB-to-SPB distance above 2 µm represents an anaphase spindle. *n*> 100 cells. (C) Maximum intensity projection of *S. cerevisiae*, tagged with GFP-Tub1 and Spc42-mCherry, and *C. albicans* tagged with Tub2-GFP and Tub4-mCherry and co-stained with Hoechst during the pre-anaphase stage.

**Figure S2.**
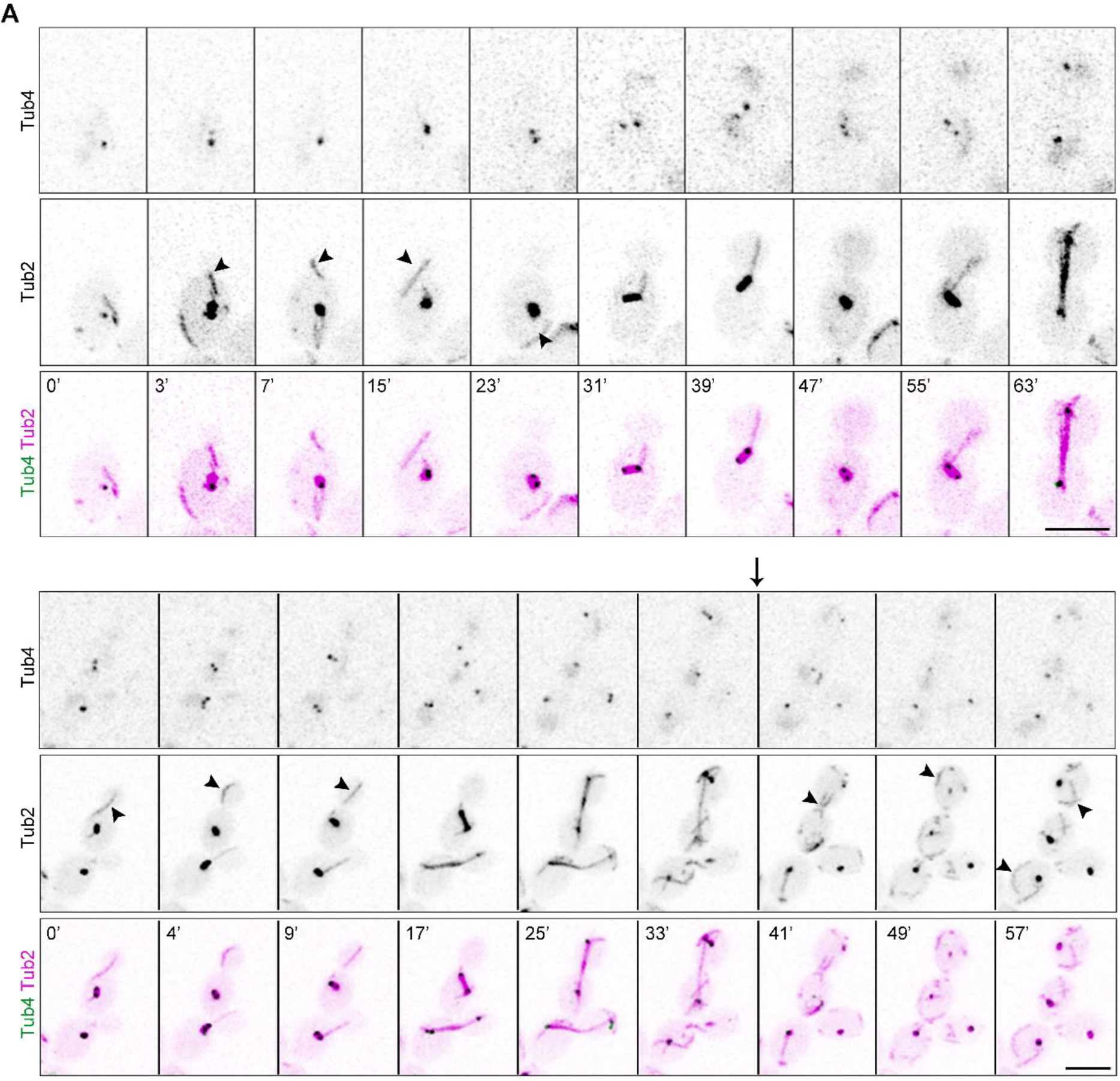
Microtubule dynamics during cell division in *C. albicans*. (A) Time-lapse images showing dynamics of the microtubule (Tub2-GFP) and SPBs (Tub4-mCherry) during the cell cycle till anaphase onset (*top*) and till completion of cytokinesis (*bottom*). The black arrowheads represent free cMTs in the cytosol. The arrow marks the onset of telophase (characterised by the disassembly of the mitotic spindle). Scale bar 5 μm.

**Supplementary Table S1:**
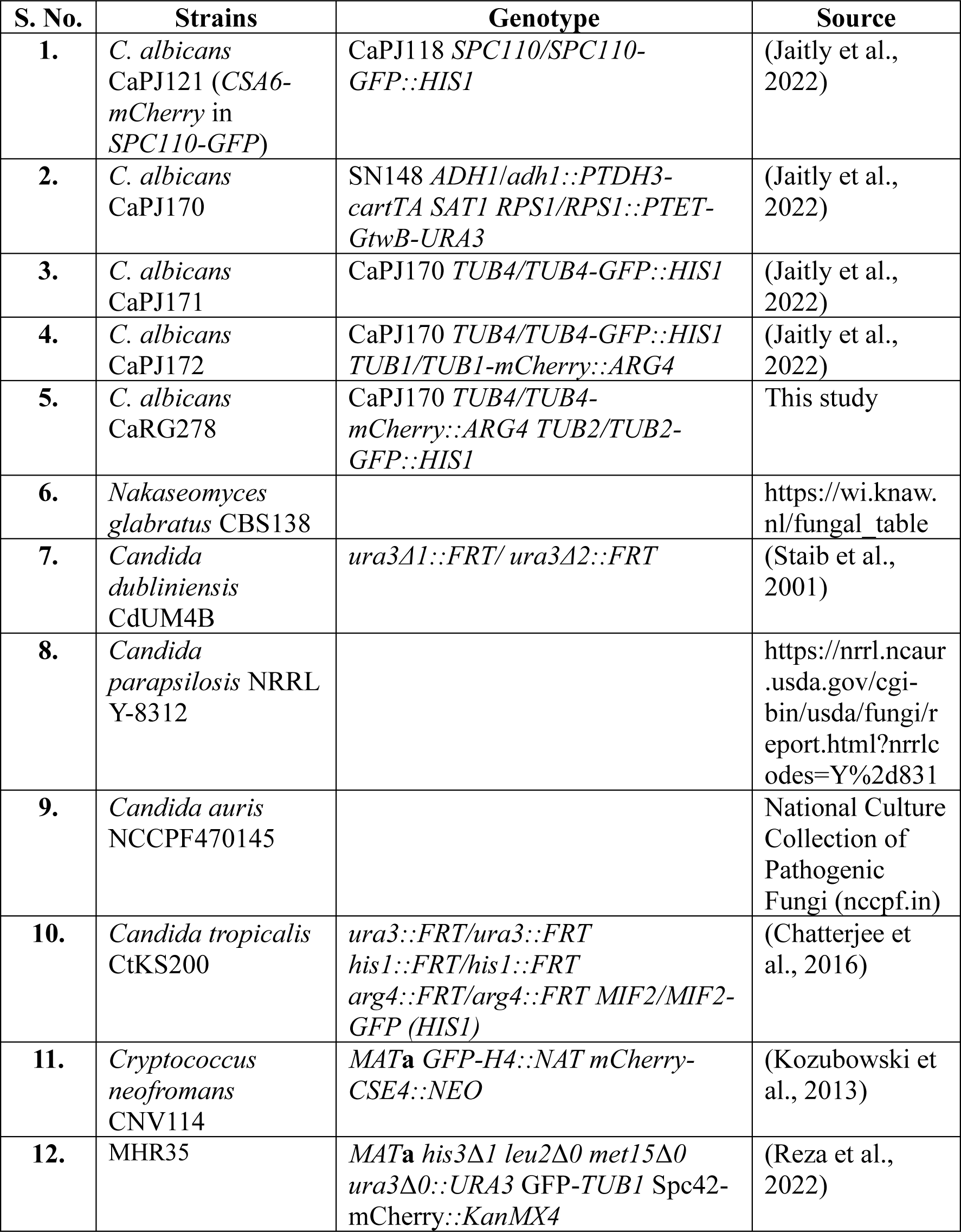
List of strains used in this study.

| S. No. | Strains | Genotype | Source |
| --- | --- | --- | --- |
| 1. | *C. albicans*<br>CaPJ121 (CSA6-<br>*mCherry* in<br>*SPC110-GFP*) | CaPJ118 *SPC110/SPC110-*<br>*GFP::HIS1* | (Jaitly et al.,<br>2022) |
| 2. | *C. albicans*<br>CaPJ170 | SN148 *ADH1/adh1::PTDH3-*<br>*cartTA SAT1 RPS1/RPS1::PTET-*<br>*GtwB-URA3* | (Jaitly et al.,<br>2022) |
| 3. | *C. albicans*<br>CaPJ171 | CaPJ170 *TUB4/TUB4-GFP::HIS1* | (Jaitly et al.,<br>2022) |
| 4. | *C. albicans*<br>CaPJ172 | CaPJ170 *TUB4/TUB4-GFP::HIS1*<br>*TUB1/TUB1-mCherry::ARG4* | (Jaitly et al.,<br>2022) |
| 5. | *C. albicans*<br>CaRG278 | CaPJ170 *TUB4/TUB4-*<br>*mCherry::ARG4 TUB2/TUB2-*<br>*GFP::HIS1* | This study |
| 6. | *Nakaseomyces*<br>*glabratus* CBS138 |  | https://wi.knaw.<br>nl/fungal_table |
| 7. | *Candida*<br>*dubliniensis*<br>CdUM4B | *ura3Δ1::FRT/ ura3Δ2::FRT* | (Staib et al.,<br>2001) |
| 8. | *Candida*<br>*parapsilosis* NRRL<br>Y-8312 |  | [https://nrrl.ncaur.<br>.usda.gov/cgi-bin/usda/fungi/r<br>eport.html?nrrlc<br>odes=Y%2d831](https://nrrl.ncaur.usda.gov/cgi-bin/usda/fungi/report.html?nrrlcodes=Y%2d831) |
| 9. | *Candida auris*<br>NCCPF470145 |  | National Culture<br>Collection of<br>Pathogenic<br>Fungi (nccpf.in) |
| 10. | *Candida tropicalis*<br>CtKS200 | *ura3::FRT/ura3::FRT*<br>*his1::FRT/his1::FRT*<br>*arg4::FRT/arg4::FRT MIF2/MIF2-*<br>*GFP (HIS1)* | (Chatterjee et<br>al., 2016) |
| 11. | *Cryptococcus*<br>*neofromans*<br>CNV114 | *MATa GFP-H4::NAT mCherry-*<br>*CSE4::NEO* | (Kozubowski et<br>al., 2013) |
| 12. | MHR35 | *MATa his3Δ1 leu2Δ0 met15Δ0*<br>*ura3Δ0::URA3 GFP-TUB1 Spc42-*<br>*mCherry::KanMX4* | (Reza et al.,<br>2022) |

